# Twitching in sleeping premature infants provides a sensitive behavioral assay of early motor control

**DOI:** 10.64898/2026.01.15.699697

**Authors:** Greta Sokoloff, Taylor G. Christiansen, Hailey C. Long, Lydia K. Karr, Olivia K. Kopp, Delaney N. Kriz, Anna C. Sanderson, Winni Liu, Isabel K. Johnson, Brenda J. Coulter, Danielle R. Rios, Tarah T. Colaizy, Mark S. Blumberg

## Abstract

Limb twitching is among the earliest observable behaviors in human development and a hallmark of active (REM) sleep. Systematic assessments in full-term infants have revealed functional features that informal observation cannot capture. Because preterm infants spend even more time asleep and are at heightened risk for neurodevelopmental disorders, we provide the first systematic characterization of twitching at 34–35 weeks postmenstrual age. Preterm infants exhibit an immense quantity of twitching across the body, underscoring its developmental significance. The spatiotemporal structure of twitching also changes with age, including a selective increase in finger and toe twitching. Unexpectedly, during periods of tracé alternant, a precursor to quiet sleep, twitching appears in brief bouts that are almost exclusively restricted to the legs. These findings show how this abundant but overlooked sleep behavior provides a sensitive assay of the developing neural control of movement, with implications for understanding typical and atypical development.

---

Two facts about active (REM) sleep capture its mystery: It predominates in early life, and it comprises a suite of components that emerge and coalesce over development [1]. One such component is myoclonic twitching: spontaneous, discrete movements of the limbs, face, and eyes. In rats, twitching begins before birth and continues into postnatal life [2]. The abundance of twitching and its diverse and unique contributions to early brain activity give rise to varied contributions to sensorimotor development [3]. Because twitching in human infants has received little formal attention among basic and clinical sleep scientists, we initiated an effort to describe its expression in full-term infants over the first six postnatal months. That effort documented many similarities and some distinct differences in the quantity and patterning of twitching between human infants and newborn rats [4,5].

Here, we turn our attention to preterm infants, for several reasons. First, the immediate postnatal period of rats corresponds most closely to the third trimester of humans [6], when active sleep (AS) is even more prevalent than it is in term newborns [7–10]. With such high rates of AS, it is not surprising that movements in fetuses [11–13] and preterm infants [8,14,15] also occur at high rates. Also, similar to newborn rats [16,17], twitches in preterm infants trigger brief cortical oscillations [18,19] that are implicated in activity-dependent brain development [3,20–23]. Thus, we have proposed that precise analysis of twitching and its neural correlates opens new avenues to understanding typical and atypical development in human infants [24,25].

Second, regarding atypical development, preterm infants are at increased risk for deficits in motor skills [26], sensory processing [27], and functional connectivity [28,29]. They are also at increased risk for cognitive delays [30] and neurodevelopmental disorders such as cerebral palsy and autism spectrum disorder [31–34]. By considering twitching within its developmental context, new opportunities arise for understanding the links between neurodevelopmental disorders and sensorimotor dysfunction [35–39], and the mediating role of sleep [40–42].

Finally, twitching in preterm infants can inform lingering issues surrounding the relative contributions of cortical and subcortical structures to early motor control. These issues linger in part due to evidence that cortical motor outflow in human infants only begins to influence behavior 3-6 months after birth [43,44]. Although brainstem motor structures, including the red nucleus (RN), are implicated in the production of twitching in infant rats and adult cats [45–47], in humans the part of the RN that produces movement is often designated as rudimentary or vestigial [48–50]. These and other factors reveal a paradox at the heart of early motor control that an assessment of twitching in premature infants can help to resolve.

The above considerations motivate our effort here to provide the first systematic analysis of the spatiotemporal structure of twitching in extremely and moderately premature human infants. We focus on infants at 34-35 weeks postmenstrual age (PMA) when cortical motor outflow is almost certainly absent [43,44]. We show that the rate and spatiotemporal patterning of twitching is similar in the extremely and moderately preterm infants, suggesting that twitching at this age is unaffected by the amount of time since birth. We show that twitching in preterm infants is abundant and dispersed across the body, and that changes in the spatiotemporal structure of twitching are developmentally continuous across preterm and full-term infants. Further, our approach demonstrates how attention to a previously overlooked sleep behavior can refine current models of brainstem involvement in early motor behavior.

## Results

### Twitching is abundant in premature infants

We recorded sleep-wake behavior during the day in extremely preterm (EPT; n=16; 8 female), moderately preterm (MPT; n=13; 5 female), and full-term (FT; n=12; 6 female) infants in this study. Premature infants were recorded at 34-35 wks PMA: Because the preterm infants were recorded at the same postmenstrual age, the chronological ages (i.e., time since birth) of the EPT infants (mean: 5.7 wks; range: 2.9-11.7 wks) were longer than those for the MPT infants (mean: 1.1 wks; range: 0.4-2.7 wks). The FT infants were recorded at 6.0 wks mean chronological age (range: 3.0-8.4 wks).

Recording sessions lasted an average of 106.9 min (range: 34.0-138.2 min) and yielded mean total sleep durations of 41.2 min for the EPT infants (range: 16.0-69.5 min), 59.2 min for the MPT infants (range: 12.8-99.6 min), and 33.6 min for the FT infants (range: 10.9-85.4 min). The EPT infants slept an average of 40% (range: 11-82%) during the recording session compared with 57% (range: 15-85%) for the MPT infants (*t*(27)=2.3, *P*<.05). Full-term infants slept an average of 64% (range: 28-98%). See Table S1 for additional participant details.

A representative 13-min recording from one MPT infant, born at 34.4 wks PMA and recorded at 35.1 wks PMA, is shown in Figure 1. During AS, twitches of the arms (shoulders/elbows, wrists, and fingers), legs (hips/knees, ankles, and toes), and face co-occur often—though not always—with rapid eye movements (REMs). Video S1 presents a 4-min segment of twitching in the same infant, accompanied by the irregular breathing that is also a hallmark of AS, followed by a transition to the high-amplitude, coordinated movements of wake. The immensity of twitching over representative 15-min sleep-wake periods across all body parts for each infant in this study is illustrated in Figure S1.

**Fig. 1.**
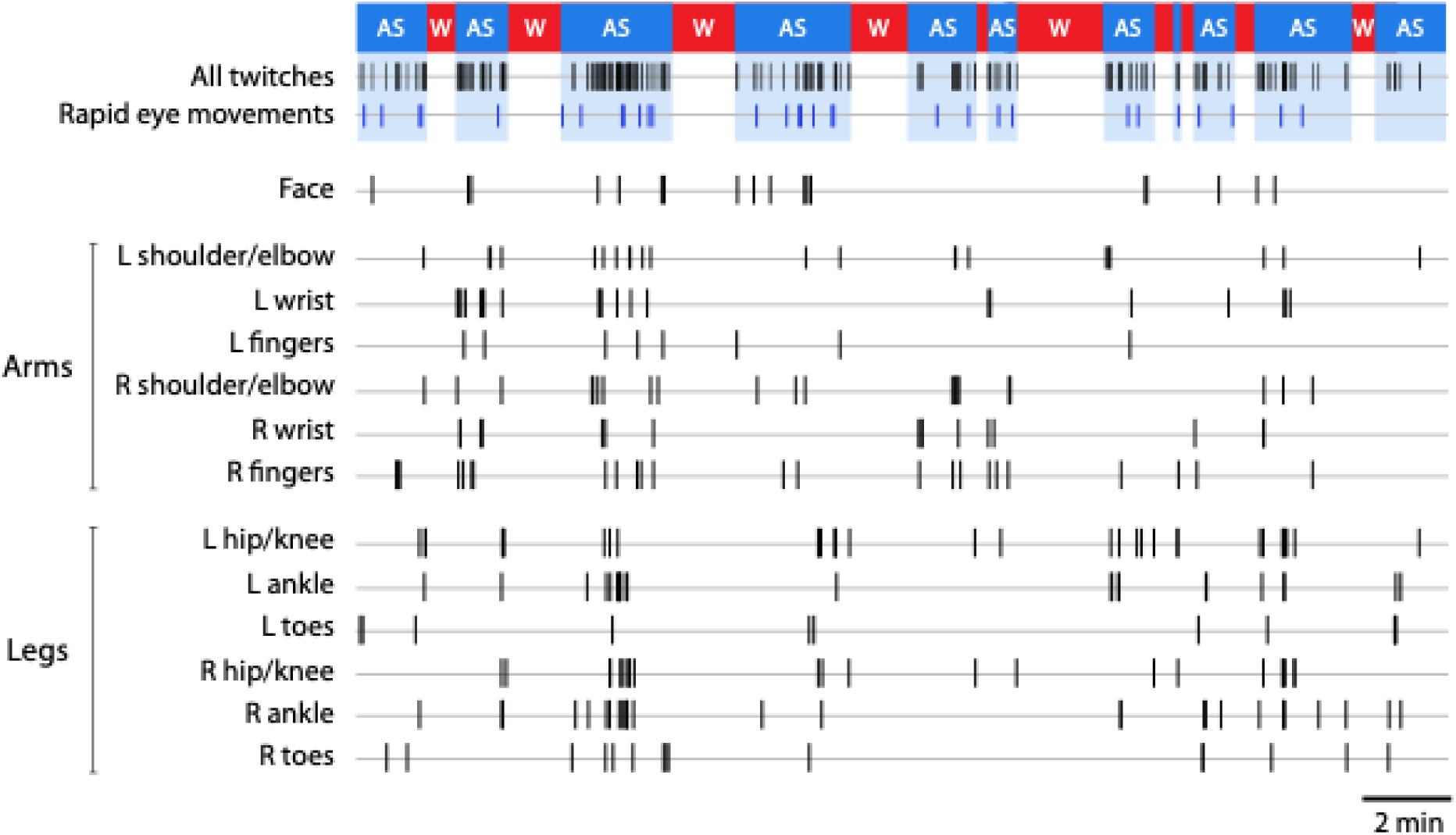
Representative data showing twitching across all body parts during active sleep in a moderately premature infant. The infant is a male born at 34.4 weeks PMA and recorded at 35.1 weeks PMA. From top to bottom: Designation of behavioral state as active sleep (AS, blue) and wakefulness (W, red). Next, all face, arm, and leg twitches are denoted as vertical ticks, as are rapid eye movements (blue ticks). Finally, twitching in all individual body parts, including left (L) and right (R) parts of the arms and legs, are shown.

Figure 2a shows the total number of scored twitches in relation to total sleep duration for each recording session for the EPT, MPT, and FT infants. The log-survivor plots in Figure 2b show that the sleep bouts of the EPT and MPT infants are similar and highly fragmented (see Figure S2), with median bout durations ranging from 54 to 82 s. The linear distributions of the bout durations, which mirror those reported in infant rats [51], signify that they occur randomly. Although the sleep bouts of the FT infants are somewhat more consolidated than those of the preterm infants, due to a subset of relatively long sleep bouts (χ*^2^*(2)=31.0, P<.0001), the median sleep-bout durations do not differ significantly among the three groups (χ*^2^*(2)=2.5; Fig. 2b, inset). Finally, it should be stressed that the sleep records were scored continuously; had we instead used epochs (or 30 s or more) to score the records, an approach that is widely used but that obscures the occurrence of short sleep bouts, the sleep-bout durations reported here would have been substantially longer.

**Fig. 2.**
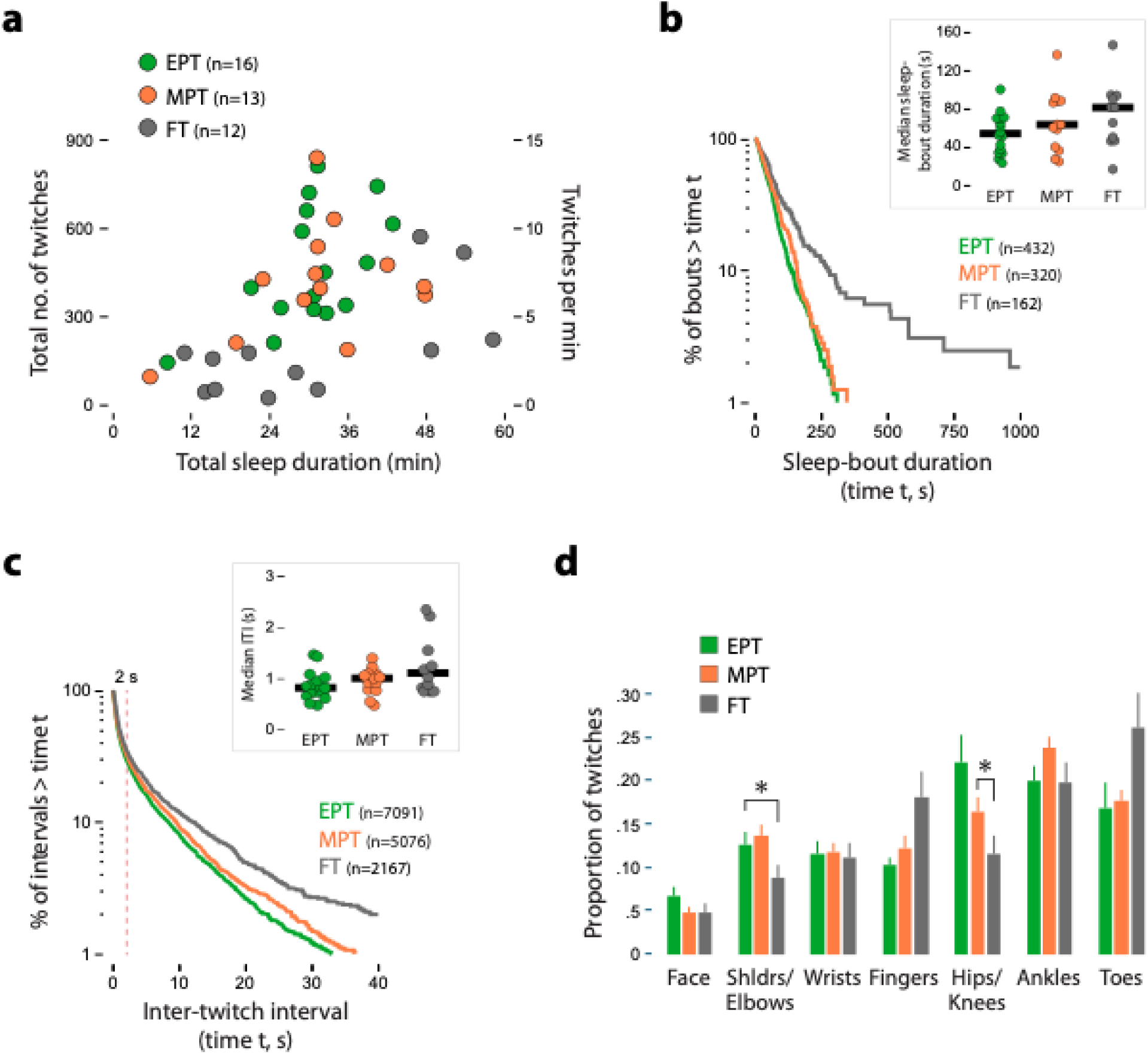
Spatiotemporal properties of sleep durations and twitching in premature and full-term infants. **a,** Scatterplot showing the total number of twitches (left axis) against total sleep duration for all the recording sessions in this study. Data are shown for extremely premature (EPT, green), moderately premature (MPT, orange), and full-term (FT, grey) infants. The right axis shows the rate of twitching (twitches/min). **b,** Log-survivor distributions of sleep-bout durations for EPT, MPT, and FT infants. Bout durations are pooled within each group. Inset: Median bout durations for each individual infant within each group, and the horizontal black bars indicate group medians. **c,** Log-survivor distributions of inter-twitch intervals for EPT, MPT, and FT infants during sleep. Intervals are pooled for each group. The vertical dashed red line shows that approximately two-thirds of ITIs are less than 2 s. Inset: Median bout durations for each individual infant for each group, and the horizontal black bars indicate group medians. **d,** Proportion of twitches during sleep across body parts for EPT, MPT, and FT infants. * significant difference between designated groups. Outlier data from one FT infant were removed from the insets in **b** and **c.**

Log-survivor plots of inter-twitch intervals (ITIs) during sleep allow us to assess their temporal structure. As shown in Figure 2c, approximately two-thirds of twitches in the EPT, MPT, and FT infants occur in bouts at intervals less than 2 s, similar to the structure observed at later ages [4]. Also, although the ITIs distribute similarly for the EPT and MPT infants, the distribution for the FT infants separates from the other two at ITIs greater than 2 s, indicating a slowing of the rate of twitching at those longer intervals. Although median inter-twitch intervals are not significantly different for the three groups (χ*^2^*(2)=4.2; Fig 2c, inset), the log-survivor distributions are significantly different (χ*^2^*(2)=66.9, P<.0001).

### The distribution of twitching across the body exhibits a developmental shift

We next assessed changes in the distribution of twitching across the body in the EPT, MPT, and FT infants (Fig. 2d). A repeated-measures ANOVA revealed a significant body part x group interaction (*F*(8.4, 160.0)=3.1, *P*<.003, adj. η^2^_p_=.10; both main effects were also significant, *P*s<.02). The interaction appears to be driven largely by a shift in twitching from the shoulders and elbows to the fingers, and from the hips and knees to the toes, although only two of the posthoc comparisons are significant (Fig. 2d).

One notable pattern in Figure 2d is the dominance of leg twitching in all three groups: the mean percentage of leg twitching relative to arm twitching is 60.6% in the EPT infants (range: 42.3-78.3%; 14/16 infants: W=1, *P*<.001); 61.0% in the MPT infants (range: 47.9-78.3%; 12/13 infants: W=7, *P*<.001), and 60.1% in the FT infants (range: 41.3-79.2%; 10/12 infants: W=13, *P*<.05).

Because there were no significant differences in the EPT and MPT groups, we collapsed them into a single preterm group and repeated the analysis (Fig. S3); again, a repeated-measures ANOVA revealed a significant body part x group interaction (*F*(4.3, 166.7)=5.0, *P*<.001, adj. η^2^_p_=.09; both main effects were again significant, *P*s<.003). Now, in the preterm infants compared with the full-term infants, we find a higher proportion of finger and toe twitching and a lower proportion of shoulders/elbows and hips/knees twitching, with no differences for the wrists and ankles.

In summary, twitching reveals the developmental onset of distal motor control in the period between 34-35 wks PMA and 8 wks after full-term birth. This shift toward distal control of the fingers and toes continues through at least five months of age [5]. Finally, this distal shift does not appear to result from extrauterine experience as the chronological age of the EPT infants was ∼4 wks greater than the MPT infants, similar to the FT infants (Table S1).

### The temporal structure of twitching reflects an emerging focus on the distal limbs

To better understand the temporal structure of twitching across the body, we performed hierarchical cluster analysis with seriation [52]. Cluster analysis enables us to assess the degree to which the twitching of body parts group (i.e., cluster) together in time. This analysis was performed on 10-s windows localized to periods of sleep, as described previously for full-term infants over the first six postnatal months [4].

The hierarchical relations among the body parts are revealed by the dendrograms at the top of Figure 3. The EPT and MPT groups exhibit similar primary structures highlighted by two clusters, one for the arms and one for the legs (the face clusters with the arms in the preterm and full-term infants). Within the arms, a secondary structure emerges characterized by (a) clustering of wrist and finger twitching within the right and left hands, and (b) clustering of shoulder/elbow twitching across the left and right arms. Similarly, within the legs, the structure is characterized by (a) clustering of ankle and toe twitching within the right and left feet, and (b) clustering of twitching in the hips/knees across the left and right legs.

**Fig. 3.**
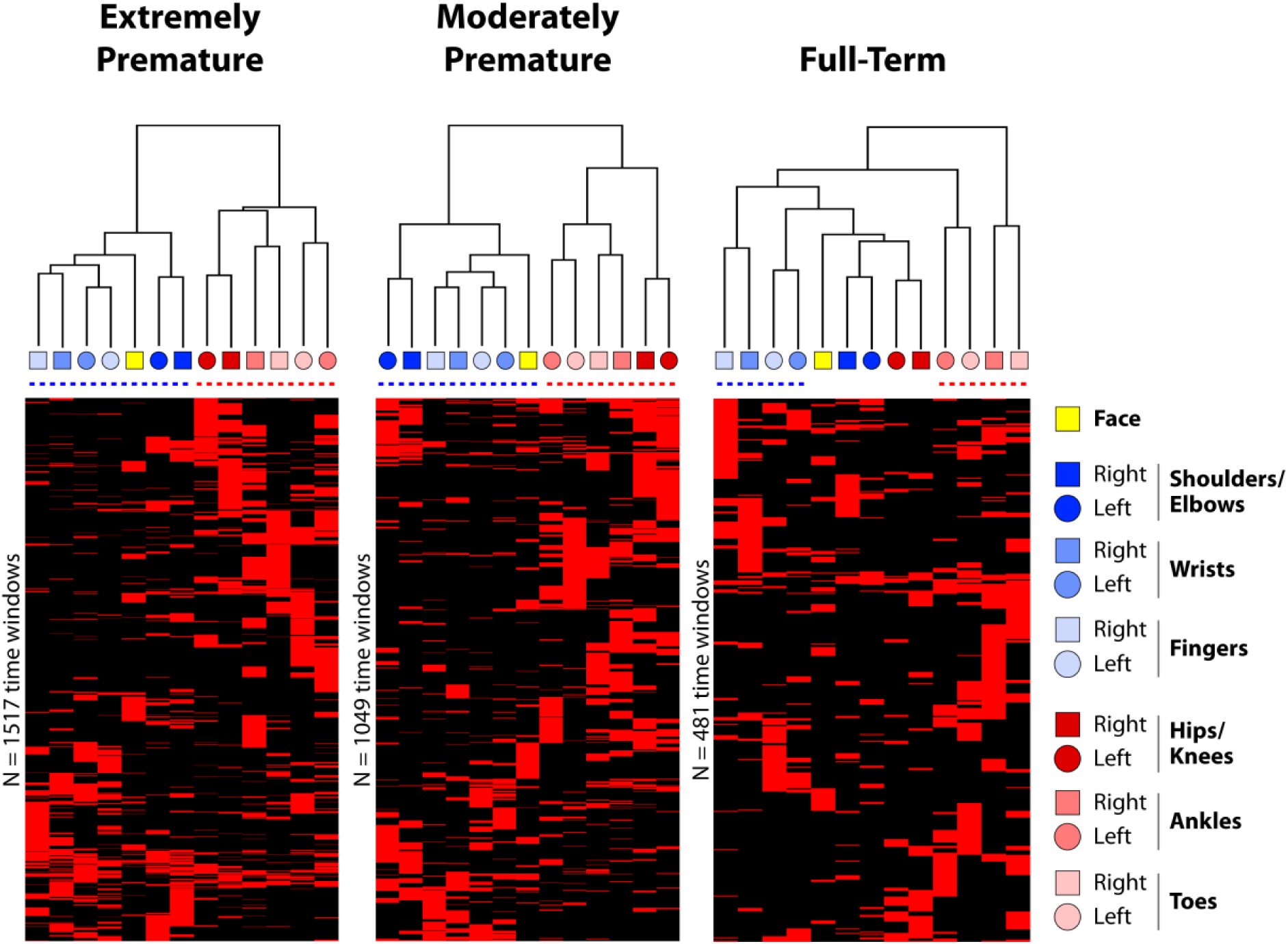
Hierarchical cluster analyses of twitching across body parts in preterm and full-term infants. Cluster analyses were performed on pooled data for extremely premature (left), moderately premature (middle), and full-term (right) infants. The key at right shows each of the body parts scored in each infant. The horizontal dashed lines beneath each dendrogram highlights the strongest clusters for the arms (blue) and legs (red). Analyses were performed on twitch data based on the presence or absence of a twitch during sequential 10-s windows during sleep. Each window is represented as a row in the cluster diagram below, with red denoted the presence of a twitch and black denoting its absence. The total number of rows are shown to the left of each cluster diagram; seriation was used to reorder the rows such that windows with similar patterns are close to each other.

Whereas the clusters in the EPT and MPT groups clearly distinguish the arms from the legs, the clusters for the FT infants are no longer segregated in that way; instead, the hierarchy is flatter, suggesting greater refinement of the control of parts within each limb, particularly the distal parts (e.g., the wrist-and-fingers and ankle-and-toes on the right and left sides of the body). Again, this shift in patterning toward the distal limbs becomes even more pronounced over the next several months in full-term infants [4] and, as with the distribution of twitching across the body, appears to be unaffected by chronological age.

### Twitching occurs unexpectedly during periods of tracé alternant

Although 50-60% of sleep in premature infants at 34-35 weeks PMA is composed of AS [8], the remainder is quiet sleep that is primarily composed of discontinuous EEG, alternating bursts of high-amplitude activity separated by inter-burst periods of low-amplitude activity [53]. Such periods begin with a discontinuous pattern of EEG bursts and subsequently transition to a continuous, periodic pattern—called tracé alternant (TA)—around the ages tested here. We identified periods of recurring EEG bursts in the EPT and MPT infants that are most consistent with TA and are hereafter designated as such. Representative data from an EPT infant, born at 26.6 weeks PMA and recorded at 34.4 weeks PMA, show several TA periods (Fig. 4a). The high-amplitude, mixed-frequency EEG bursts (Fig. 4b) resemble those reported previously [54].

**Fig. 4.**
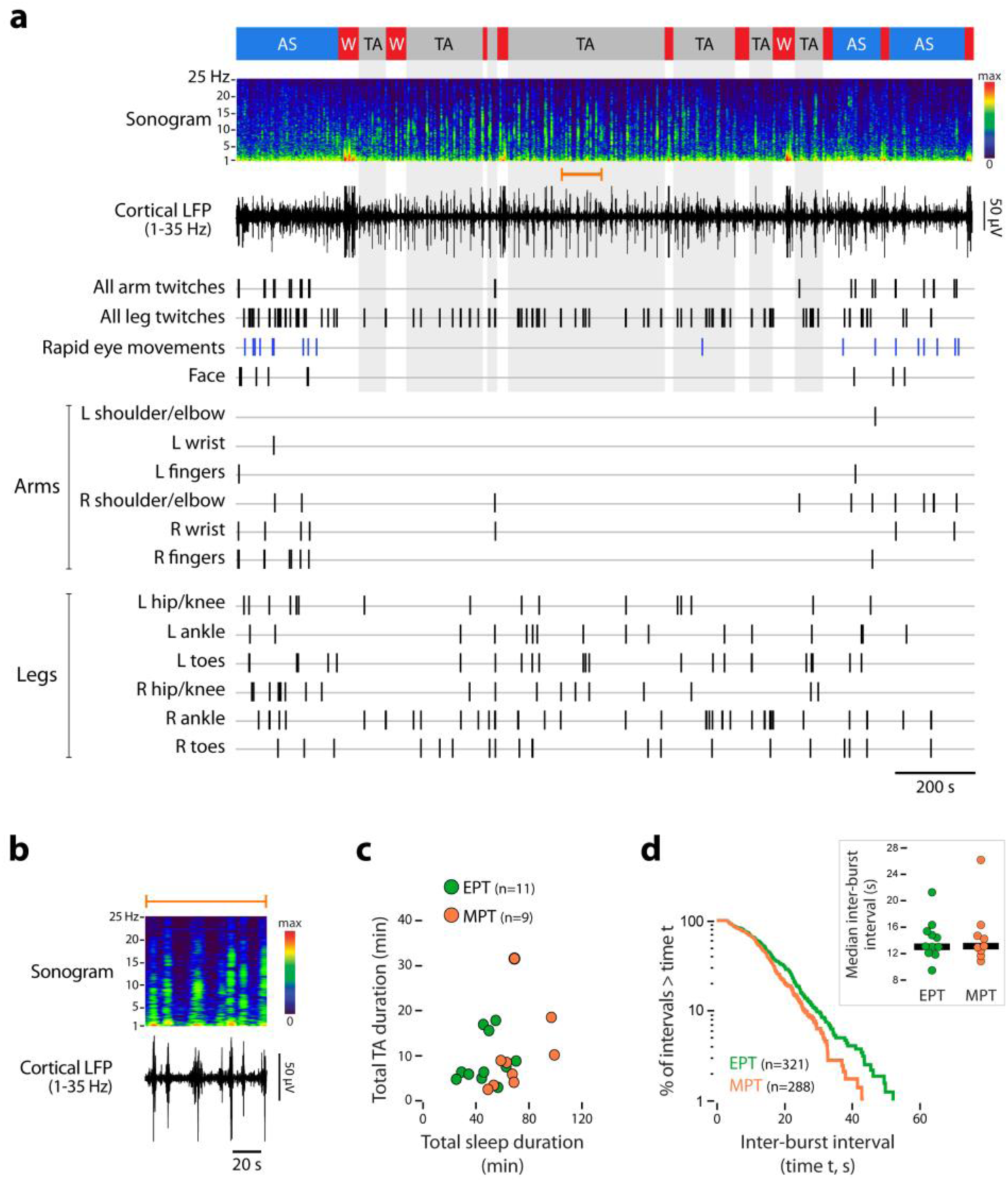
Twitching occurs during periods of tracé alternant. **a,** Representative data from an extremely premature male infant born at 26.6 weeks PMA and recorded at 34.4 weeks PMA. From top to bottom: Designation of behavioral state as active sleep (AS, blue), tracé alternant (TA, grey) and wakefulness (W, red), followed by cortical local field potential (LFP; 1-35 Hz) and associated sonogram. Next, all arm twitches and leg twitches are denoted as vertical ticks, as are rapid eye movements (blue ticks). Finally, twitching in all individual body parts, including left (L) and right (R) parts of the arms and legs, are shown. **b,** Enlarged section of cortical LFP and associated sonogram for the orange-bracketed segment in **a**. **c,** Scatterplot showing the total duration of TA periods against total sleep duration for the extremely premature (EPT, green) and moderately premature (MPT, orange) infants. **d,** Log-survivor distributions of the intervals between high-amplitude EEG bursts during TA periods for EPT and MPT infants. Intervals are pooled within each group. Inset: Median inter-burst intervals for each individual infant within each group, and the horizontal black bars indicate group medians.

Overall, 11 of the EPT infants and 9 of the MPT infants exhibited clear TA periods, with a mean total duration of 9.3 min (range: 2.4-31.3 min; Fig. 4c). The EEG bursts occurred rhythmically at median intervals of ∼13 s (Fig. 4d); although the log-survivor distributions of inter-TA intervals indicated a small but significant difference between the EPT and MPT infants driven by deviations at the longer intervals (χ*^2^*(2)=5.78, P<.05), the median intervals were not significantly different (*U*=47).

TA is conventionally defined as a quiet-sleep-like period characterized by regular respiration and an absence of REMs and twitching. Surprisingly, however, we found that twitching does occur during this state, as illustrated in the representative data from an EPT infant in Figure 4a (see also Video S2). Careful visual inspection of the movements indicated that they are indistinguishable from twitches during AS, though they represent only ∼10% of all twitches produced in those 20 premature infants that exhibited periods of both AS and TA. The temporal distributions of inter-twitch intervals during AS and TA (collapsed across EPT and MPT infants) are identical up to ITIs of ∼800 ms, after which the curves diverge significantly (Fig. 5a; χ*^2^*(1)=91.8, P<.0001). The sharp inflection point at ∼800 ms (Fig. 5a, inset) indicates that twitching during TA tends to occur in more concentrated bursts than does twitching during AS.

**Fig. 5.**
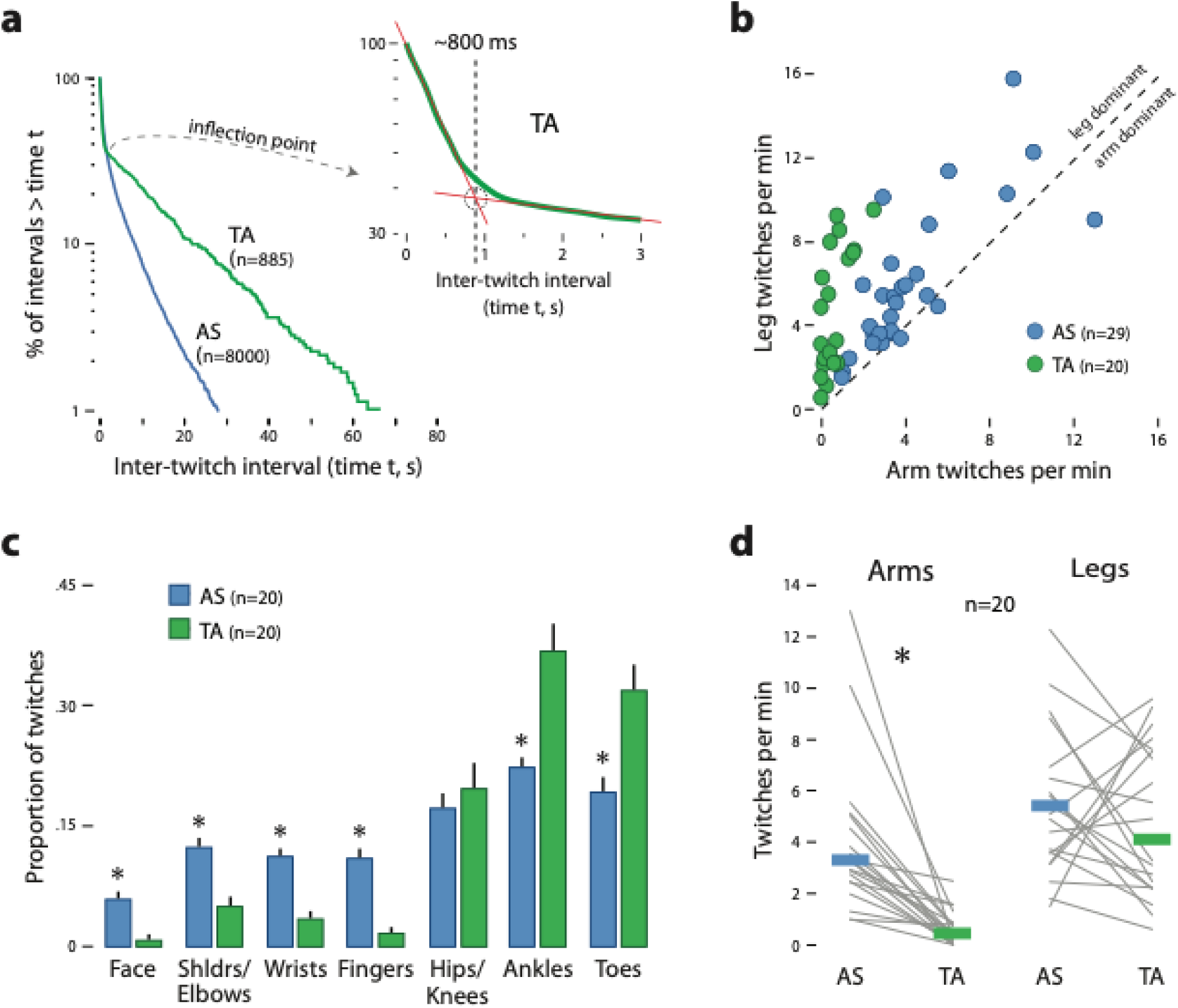
Spatiotemporal features of twitching during periods of tracé alternant in premature infants. **a,** Log-survivor distributions of inter-twitch intervals for premature infants during active sleep (AS, blue) and periods of tracé alternant (TA, green). Intervals are pooled for each group. Inset: Magnification of the TA plot at left to show the distributions where the TA and AS plots diverge; the inflection point indicates that bouts of twitching during TA periods do not last longer than ∼800 ms. **b,** Scatterplot showing the rate of leg twitching versus the rate of arm twitching for all infants with periods of AS and/or TA. Points above the line of isometry (dashed diagonal line) indicate infants with higher rates of leg twitching. **c,** Proportion of twitches during sleep across body parts for infants exhibiting periods of both AS and TA. * significant difference from other group. **d,** Median twitch rate for the arms (left) and legs (right) during periods of AS (blue bars) and TA (green bars) in infants that exhibited both states. Paired data for individual infants are shown as grey lines. * significant difference between the two groups.

We next assessed whether the EEG bursts and twitching during TA are temporally linked but were unable to find any evidence of such a linkage. The rate of twitching during the periods encompassing EEG bursts (4.5 ± 1.0 twitches/min) and those between EEG bursts (5.6 ± 0.8 twitches/min) did not differ significantly (*t*(19)=0.9), nor were there differences between the EPT and MPT infants (*t*(18)=0.0). Therefore, unlike the association between twitches and sleep spindles during quiet sleep in full-term infants beginning around 3 months of age [5,55], there appears to be no association between twitches and EEG bursts during TA periods.

The bodily distribution of twitches during TA exhibits a particularly striking pattern, namely, an extreme bias toward the legs (Fig. 5b). On average, leg twitching during TA accounts for 91% of all limb twitches (legs>arms in 20/20 infants: *W*=0.0, *P*<.001) compared with 61% during AS (legs>arms in 26/29 infants: *W*=26.0, *P*<.001). Looking across body parts, the leg bias during TA was expressed in the ankles and toes, not the hips/knees (Fig. 5c). Overall, a repeated-measures ANOVA (necessarily restricted to only those 20 infants that exhibited periods of both AS and TA) revealed a significant body part x state interaction (*F*(4.7, 89.4)=25.5, *P*<.001, adj. η^2^_p_=.55; the main effects were also significant). Finally, whereas 20/20 infants exhibited higher rates of twitching in the arms during AS than during TA (W=253, *P*<.001), for the legs that was true for only 13/20 infants (W=156, *NS*; Fig. 5d). Altogether, the unique spatiotemporal structure and extreme leg bias of twitches during TA suggests that they are phenomenologically distinct from twitches during AS.

## Discussion

In common parlance, “to sleep like a baby” is to sleep deeply and well. Ironically, the sleep of actual babies looks very different, featuring many short bouts of sleep interspersed with equally short awakenings. In addition, infant sleep often appears agitated due to the presence of limb and facial twitches, rapid eye movements (REMs), and irregular breathing, which are characteristics of AS, the predominant behavioral state of early life. Although previous studies have documented the presence of spontaneous movements in fetuses and preterm infants, this is the first study to explore the spatiotemporal properties of twitching in premature infants. Our findings parallel those in infant rats [56] by showing that twitching is abundant in human infants and its structure becomes increasingly more organized with age. Moreover, our findings suggest that twitching provides a unique and powerful behavioral assay of the developing neural representation of the limbs.

Our findings also provide evidence for developmental continuity with regard to some aspects of the expression of twitching. The notion of developmental continuity is complex [57,58], but here we use the concept to refer to similarities in several core features of twitching in premature and full-term infants. Those core features include the discrete, jerky nature of the movements, their occurrence during sleep, their marked contrast with wake movements, and their spatiotemporal features. Combined with our earlier assessments of twitching in full-term infants over the first six postnatal months [4,5], the addition of data from premature infants provides compelling evidence of developmental continuity in the early expression of key aspects of the behavior—including gradual changes in the distributions of inter-twitch intervals (Fig. 2c) and in the proportion (Fig. 2d) and temporal clustering (Fig. 3) of twitching across body parts. We stress, however, that this assertion of continuity does not necessarily imply continuity in all aspects of twitching across these ages. Indeed, in addition to those features of twitching that appear continuous, there are other features that are unique to specific stages of life, including the expression of twitching during periods of tracé alternant in preterm infants (as found here) and during quiet sleep in full-term infants beginning around three months of age [5].

Psychologists have long noted how the quantity of experiential input—for example, how many words children hear [59] or how many steps children take [60]—play formative roles in cognitive and motor development. Similarly, we previously estimated that full-term infants produce approximately 600 twitches/h; with 8 h of AS each day, that rate yields a daily output of ∼4800 twitches [4]. With preterm infants twitching at a similar rate but with a daily quota of AS that is roughly double that of full-term infants [61], the daily quantity of twitches may approach 10,000. By providing discrete opportunities for relating self-generated motor output with sensory input, twitching contributes in multiple ways to sensorimotor plasticity [3]. The immense quantity of twitching only bolsters its developmental significance (Fig. S1).

It is notable that we did not detect any group differences between the EPT and MPT infants on the dimensions of twitching or sleep-bout organization analyzed here. Such differences might have been expected given that the EPT infants were an average of 5 wks older chronologically than the MPT infants. Therefore, that twitching does not differ between the two groups suggests that its developmental course is unaffected by extrauterine experience.

The lack of a difference in twitching between EPT and MPT infants is surprising given that EPT infants are more likely to exhibit motor impairments, including cerebral palsy [62]. However, developmental outcomes are highly variable and may not show up as group differences, especially at early ages. Instead, it may be more informative to assess sleep and twitching in premature infants with low body weight at birth and with evidence of early neurological insult. We are continuing to follow these and the other infants in this study to assess possible relations among twitching, motor development, and the emergence of neurodevelopmental disorders. Several of the findings here will be used to inform those assessments; for instance, individual differences among preterm infants in the expression of twitching across the body, the emergence of distal limb control, the occurrence of leg twitching during TA, and/or the bias toward leg movements may predict deficits in motor control.

Although we cannot directly assess the neural mechanisms that produce twitching in human infants, research in non-human animals provides valuable guidance. Specifically, in newborn rats [63] and adult cats [64] the cortex has little or no role to play in the production of twitches. Instead, brainstem motor structures, including the red nucleus (RN), are implicated in the production of twitching in these animals [45–47]. The RN has two parts: the magnocellular part—the source of the rubrospinal tract—generates motor commands, and the parvocellular part modulates cerebellar circuits but plays no direct role in motor behavior [49,50,65]. In human adults, the relative prominence of the corticospinal tract contrasted with the small volume and limited spinal connectivity of the magnocellular RN justifies the latter’s designation as a rudimentary or vestigial structure [48–50]. These observations raise an intriguing paradox: *If the magnocellular RN is vestigial in humans and cortical motor outflow is developmentally delayed, how do we explain the prominence of twitching in premature infants?*

One possible resolution of this paradox is to consider that the reticulospinal and vestibulospinal tracts, which are also implicated in the production of twitching [64,66], compensate for the absence of a functioning magnocellular RN. However, a role for the magnocellular RN cannot be ruled out because studies in human fetuses reveal that the magnocellular RN is actually larger than the parvocellular RN and continues to enlarge before birth [67,68]. The authors of one of these studies suggested that the magnocellular RN in early human development contributes to “a specific transitory pattern of motor behaviour” [68] (p. 95). Based on the present findings, we propose that the production of twitching is a major, if not primary, function of the magnocellular RN in preterm infants, and predict that the reduced rates of twitching in full-term infants across age will mirror the diminution of the magnocellular RN’s size and contribution to behavior.

Thus, humans may be similar to rats in relying on the magnocellular RN and other brainstem structures for the production of twitching, at least in early development [47,63]. If so, then the developmental changes found here in the spatiotemporal properties of twitching—including the increased focus on twitches of the distal limbs—suggest developmental tuning of motor maps in the brainstem [69].

Our findings raise other questions as well. For example, it is not clear what accounts for the 60:40 leg:arm bias in twitching in the preterm infants and the full-term neonates. This bias, which disappears by 5-6 months postnatal age [5,55], may reflect transient anatomical biases in motor projections to the cervical and lumbosacral spinal cord. That twitching occurs during TA—and that it exhibits an extreme 90:10 leg:arm bias—was a surprise given that, to our knowledge, TA-related twitching has not been previously reported. The absence of such reports is likely attributable to the widespread use of swaddling in the NICU, a practice that obstructs visual observation of the legs. The extreme leg:arm bias may reflect selective and sleep-specific activation of (for example) the lateral vestibulospinal tract whose axons project directly to the lumbar spinal cord [70,71]. Nonetheless, these biases cannot be explained at this time.

Similarly, it is difficult to account for the selective increase in finger twitching that emerges between 34-35 wks PMA and 3-8 wks postnatal age. This developmental shift need not reflect the emergence of corticospinal circuits as both the rubrospinal and reticulospinal tracts of primates are also implicated in finger movement [72,73]. Indeed, the onset of corticospinal function may be more specifically associated with independent finger movements [74], which emerge during wake behavior only after 6 months of age [75]. Independent control of the digits is an aspect of twitching that remains to be systematically assessed. Such assessments should also include the toes as they twitch at similarly high rates as the fingers in preterm infants, as well as full-term infants through at least 5-6 months of age [5,55]. Although such high rates of toe twitching may seem surprising, it should be noted that human infants reach for objects with their feet at earlier ages than they reach with their hands [76], a possible remnant of our primate heritage [77].

Given the large amount of sleep in fetuses and infants [45], one might assume that protecting sleep is a central concern of pediatric facilities—especially NICUs. However, only recently have neonatologists, noting basic sleep research, reassessed the often brightly lit, noisy, and disruptive NICU environments that preterm infants traditionally experience [78–82]. In addition to improving the external environment within the NICU, efforts are being made to implement safe sleep practices that focus on the environment within the incubator [83,84]. These efforts complement a growing appreciation for the importance of sleep, including AS [40,85], to short- and long-term development. Nonetheless, as a prominent spontaneous behavior and core component of AS, twitching has been variously dismissed in adults as a by-product of dreams [86] or, in fetuses and premature infants, as a sign of distress [11,87]. Instead, we encourage neonatologists and developmental scientists to reconsider this behavior in light of accumulating evidence of its heuristic value and clinical significance.

## Methods

### Participants

A total of 29 preterm infants participated in this study. Premature infants were divided into two groups: extremely preterm (EPT; <32 weeks postmenstrual age at birth; n=16; 8 female) and moderately preterm (MPT; 32-37 weeks postmenstrual age at birth; n=13; 5 female). An additional cohort of full-term infants (FT; 3.0-8.4 weeks of age at testing; n=12, 6 female) are also included.

Preterm infants were drawn from a pool of infants recruited and consented into the discovery cohort of a larger, ongoing project conducted in the Neonatal Intensive Care Unit (NICU) at the Stead Family Children’s Hospital at the University of Iowa Hospital and Clinics. Preterm infants for this study were selected based on age (i.e., 34-35 wks PMA) and the ability to visualize the arms, legs, and face in the video recordings. Full-term infants were recruited from the community using email and print advertisements and were similarly drawn from a pool of infants for the larger study. Table S1 provides demographic, age, weight, and sleep-session information for all 41 infants included in this study. Guardians identified the race of the infants as 65.9% White, 14.6% two or more races, 4.9% African American/Black, and 2.4% Asian; 12.2% preferred not to specify. Guardians also identified the ethnicity of the infants as 19.5% Hispanic or Latino; 7.3% preferred not to specify. All study procedures were approved by the Institutional Review Board at the University of Iowa.

### Recording of sleep sessions

Preterm infants consented into the study participated in multiple sessions, at approximately 2-week intervals, while in the NICU. All NICU recording sessions were conducted by clinical research nurses and took place between 0800 and 1600, preferably after a feeding. While in an incubator or crib, in a supine position, infants were fitted with two pairs of Neonatal Hydrogel Sensors connected to an Olympic Brainz Monitor (Natus Medical Incorporated, Middleton, WI). After preparing the scalp with Nuprep (Weaver and Company, Aurora, CO) and coating the electrode with conductive paste (Ten20, Weaver and Company), electrodes were secured at standard EEG sites (C3/C4, P3/P4) using neonatal positioning strips (Natus) and a surgical skin marker; an additional ground electrode was secured to the shoulder blade. Vital signs of all infants in the NICU, including respiration, heart rate, and blood-oxygen levels, were measured and stored continuously using Sickbay (Medical Informatics Corporation, Houston, TX). An iPhone was placed above the infant on the surface of the incubator or attached to an adjustable stand next to the crib to acquire video during sleep sessions. Synchronization of the aEEG, Sickbay, and video data occurred offline.

Full-term infants were recorded in a supine, semi-reclined position in the sleep laboratory, while wearing a lab-supplied onesie, as described previously [5]. Participants were prepared for the recording session by fitting them with a high-density (124-site) EEG electrode cap (Geodesic Sensor Net; Magstim EGI, Roseville, MN). Before application, the cap was soaked in an electrolyte solution composed of potassium chloride and baby shampoo (Johnson & Johnson, New Brunswick, New Jersey) dissolved in warm water. To record respiration, a piezoelectric sensor (Unimed Electrode Supplies, Surrey, UK) was taped to the onesie on the abdomen just below the xyphoid process. The infant was recorded from a frontal view using a high-resolution video camera. EEG, respiration, and video data were synchronized during acquisition.

### Sleep staging

Videos of sleep sessions were scored using Datavyu [88]. For all videos, two experienced raters visually scored 100% of the video for sleep, wake, startles, and arousals. All videos were scored continuously, that is, epochs were not used. Scorers also noted periods when the infant’s face or limbs were obscured. Sleep and wake were scored as described previously [4,5]. Briefly, we used established behavioral criteria to define sleep and wake [7]. During wake, infants’ eyes may be open for sustained periods, eye movements are horizontal and focused, postural tone is high, and the infant exhibits high-amplitude and often synchronous limb movements [89]. During sleep, the eyes are typically closed (although they may be open) and postural tone is low; during AS, myoclonic twitches, REMs, and irregular breathing are also evident [8,89]. Importantly, twitch movements differ from wake movements in that they are relatively brief, low in amplitude, and discrete.

Periods of tracé alternant (TA) were defined as consisting of a background of low-amplitude EEG interrupted at regular intervals by high-amplitude bursts (∼100 µV). All other sleep periods were designated as AS. Finally, all periods in which nurses or clinicians were providing bedside care or a guardian was interacting with the infant were noted and excluded from analysis.

### Twitch coding and reliability

Experienced behavioral coders reviewed sleep periods for myoclonic twitches using methods similar to those described previously [4,5]. Briefly, in Datavyu, coders—blind to the EEG record—marked the onset and offset times for twitches of the face, arms, legs, hands, feet, fingers, and toes. REMs were also scored. Limb twitches were not scored if they were artifacts of other movements, such as breathing, stretching, or twitching in another limb. The average twitch durations of ∼630 ms were uniform across the infants in all three groups, consistent with our finding in rats that twitches are highly stereotyped across age [90].

A primary coder was assigned to each session to score twitches during designated sleep periods. For sessions with fewer than 30 min of sleep, all sleep periods were scored; for sessions with more than 30 min of sleep, sleep periods with optimal limb visibility were selected for scoring. For each session, a secondary coder was assigned to score at least 25% of the scored periods to assess inter-rater reliability (IRR). IRR was calculated for each period scored by both coders to quantify their level of agreement regarding the onset times and durations of all twitch events (5). Periods of disagreement between coders were identified and reviewed independently by both coders. The median value of Cohen’s kappa was 0.848 (range: 0.617-0.968). A final pass with both coders resolved any remaining discrepancies by mutual agreement. Finally, only twitch-related data scored by the primary coder were used for analysis.

### Exclusion criteria

Premature infants in the NICU were not approached for consent into the study if they were on high-frequency ventilation, tested positive for COVID-19, and/or were administered opioids or benzodiazepines as part of their care plan. In addition, one FT infant had only one very long sleep bout during the session, which resulted in anomalous median values for sleep-bout duration and inter-twitch interval; those data were excluded from those two analyses.

### Data analysis

#### Total sleep time and sleep-bout durations

The duration of each sleep bout was determined. Sleep bouts were pooled across infants within groups and JMP Pro 18 (JMP Statistical Discovery, Cary, NC) was used to plot log-survivor distributions.

#### Counts and rates for twitching

Counts for twitches (of the face, arms, and legs) and REMs were summed across periods of sleep and divided by the total time asleep. For some analyses, rates of twitching were computed separately during AS and TA.

#### Inter-twitch intervals

The time between successive twitches within a sleep period was defined as an inter-twitch interval (ITI). ITIs were excluded when there was an intervening startle, microarousal, or transition between states. ITI data were pooled across infants within groups (EPT, MPT, FT), and JMP Pro 18 was used to plot log-survivor distributions.

#### Proportion of twitching in individual body parts

To assess the distribution of twitching across the face, arms (shoulders/elbows, wrists, fingers), and legs (hips/knees, ankles, toes) for each sleep session, the number of twitches in each part was divided by the total number of twitches during the session. Proportions were arc-sine transformed before statistical analysis.

#### Cluster analysis

To assess the spatiotemporal patterning of twitching, hierarchical cluster analysis with seriation was performed using PermutMatrix software [52]. The settings were Manhattan distance and Ward’s minimum variance; for seriation, we used the multiple-fragment heuristic. Using an approach similar to that described previously [4], sequential 10-s windows were created within each sleep session; each window began at the onset of the first twitch. Twitches of the face, arms, and legs were included. To ensure that clustering was not biased by the occurrence of more than one twitch of a single body part, the maximum value for each body part within a window was 1. For the cluster analysis, data were pooled within groups. In addition to producing dendrograms showing clustering among body parts, the windows (i.e., rows) were reordered using seriation to reveal their structure.

#### Cortical EEG bursts during tracé alternant (TA), TA-period durations, and inter-burst intervals

TA periods began when an alternating EEG trace was observed with bursts exceeding 50 µV and typically a 25-50% reduction in EEG amplitude during the inter-burst intervals (see Fig. 4a). TA periods ended with the cessation of bursts and an increase in baseline EEG amplitude. To capture the EEG bursts associated with TA, EEG data were downsampled to 100 Hz and low-pass filtered at 2 Hz. Filtered data were transformed by taking the root mean square (RMS, 150-ms time constant). Over entire TA periods, the average RMS amplitude of the filtered signal was calculated and then multiplied by 3.5 to create a threshold for detecting EEG bursts. The threshold was used to determine the onset of a burst and a 3-s step was added to minimize double counting. Inter-burst intervals were calculated as the time between successive bursts within the TA period. Inter-burst intervals that spanned a transition from TA to another state (e.g., wake, AS) were removed.

### Statistical analysis

Statistical analyses were conducted using JMP Pro 18 or jamovi [91]. Before conducting ANOVAs, data were tested for normality using the Shapiro-Wilks test and for sphericity using Mauchly’s test. When sphericity was violated, a Huynh-Feldt correction was applied to the degrees of freedom. The measure of effect size for ANOVAs is represented as partial eta-squared (η^2^_p_), adjusted for positive bias [92]. Group differences in log-survivor distributions were tested using the Mantel-Cox log-rank test, which gives equal weight to all time points; because data were pooled across infants within each group, these tests should be interpreted with caution. Group differences in median sleep-bout duration, median ITI, and median inter-TA interval were tested using the non-parametric Kruskal-Wallis χ^2^ test or Mann-Whitney U test, as appropriate. The Wilcoxon signed-ranks test was used to assess leg dominance within the AS and TA groups, and the Wilcoxon matched-pairs signed-ranks test was used to assess within-infant differences in the rate of arm and leg twitching. Unless otherwise noted, means are presented with standard errors, and alpha was set at 0.05.

## Reporting summary

Further information on research design is available in the Nature Portfolio Reporting Summary linked to this article.

## Data availability

Before publication, summary data, individual data, and those videos with appropriate consents will be made available on Databrary.org.

## Code availability

Before publication, the code for the analyses will be made available on Github.

---

**Fig. S1.**
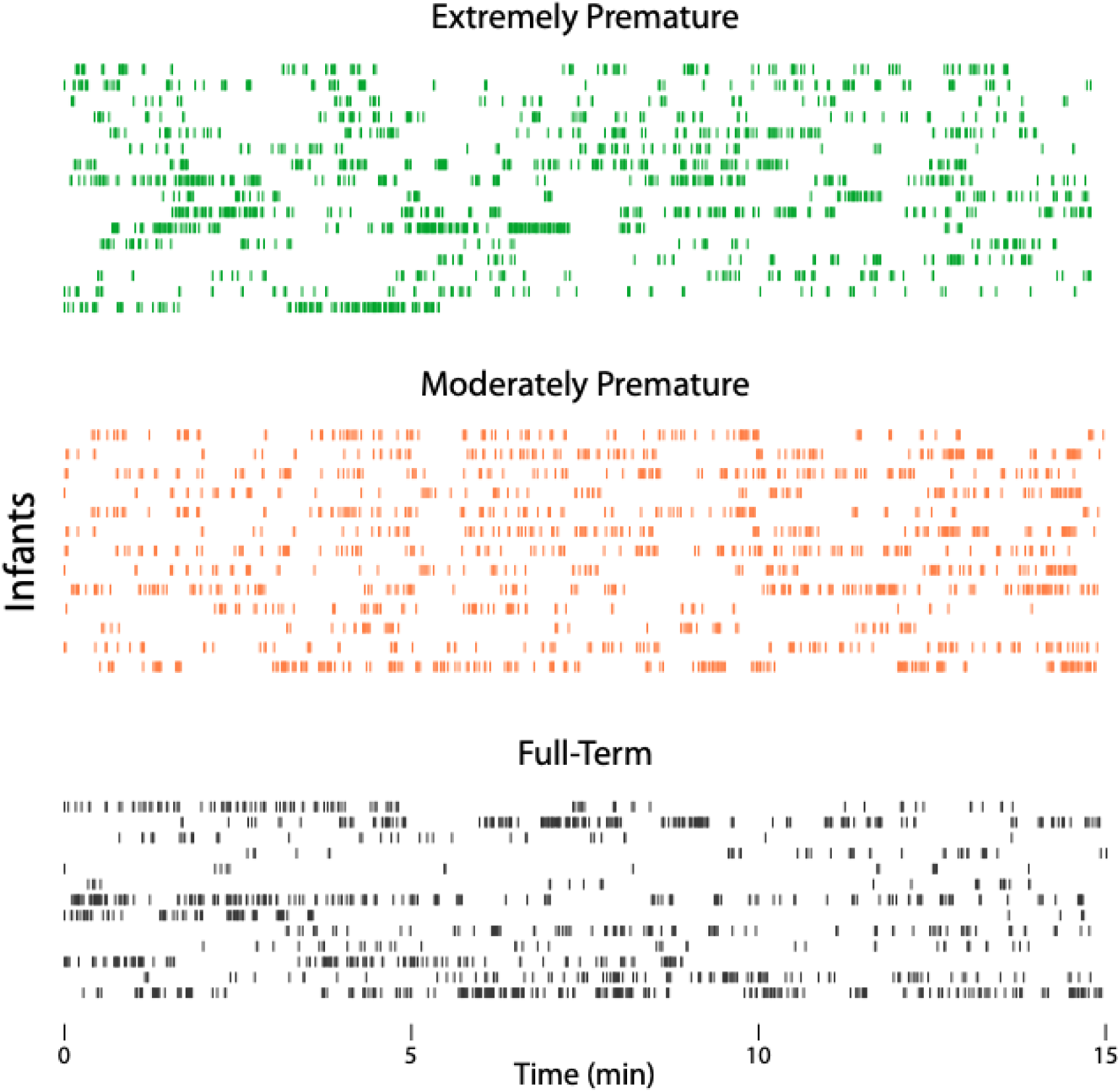
Raster plots showing individual twitches for the extremely premature (green), moderately premature (orange), and full-term (grey) infants. Each row represents an individual infant across sleep and wake periods (behavioral states are not indicated). Only 15 min of twitching are shown from each recording session.

**Fig. S2.**
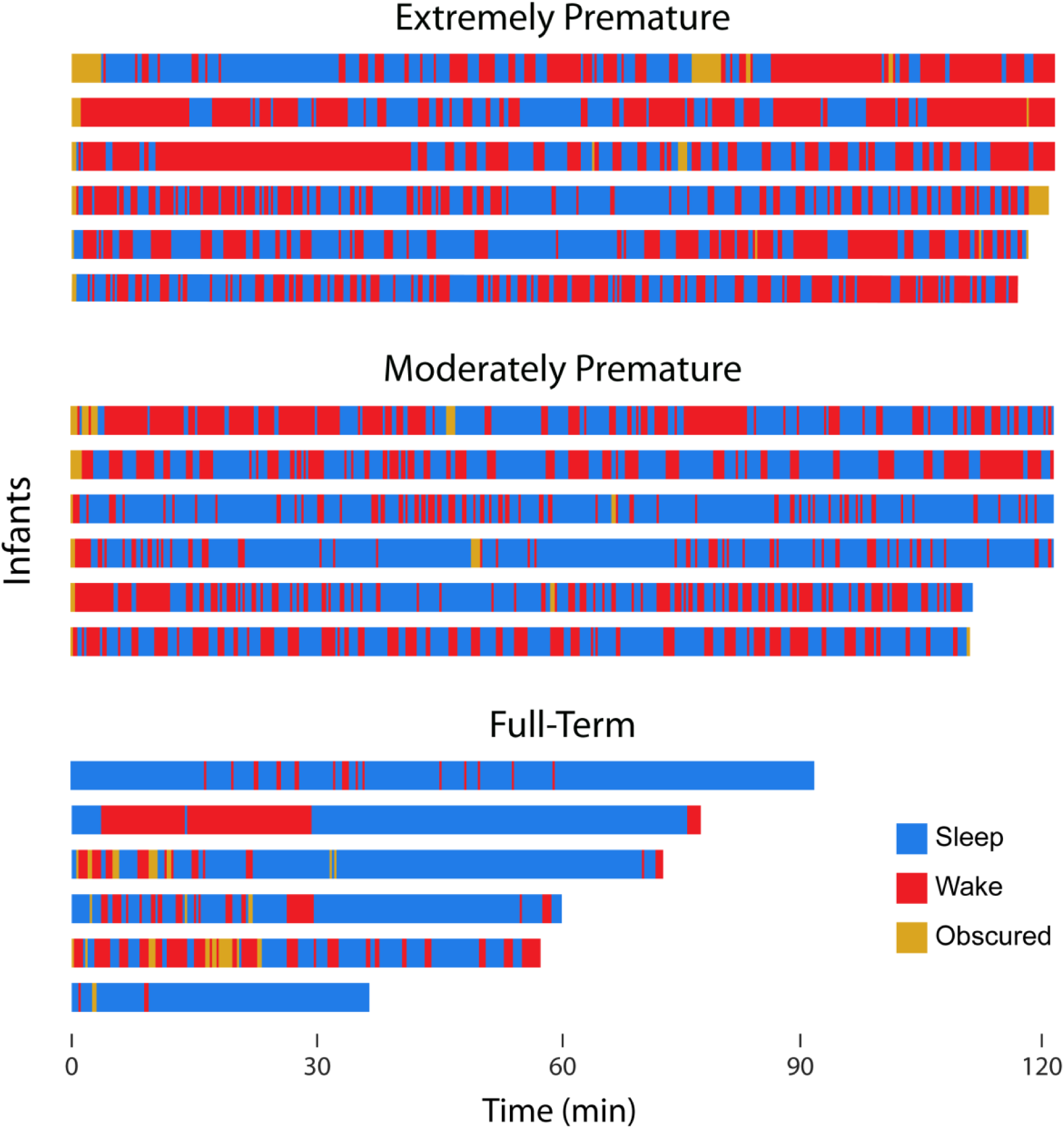
Raster plots showing periods of sleep (blue) and wake (red) for subsets of extremely premature, moderately premature, and full-term infants. Periods when the infant was obscured from view are also shown (gold). Five infants from each group are shown, with each infant represented by a different row. Records are sorted from longest to shortest within each group.

**Fig. S3.**
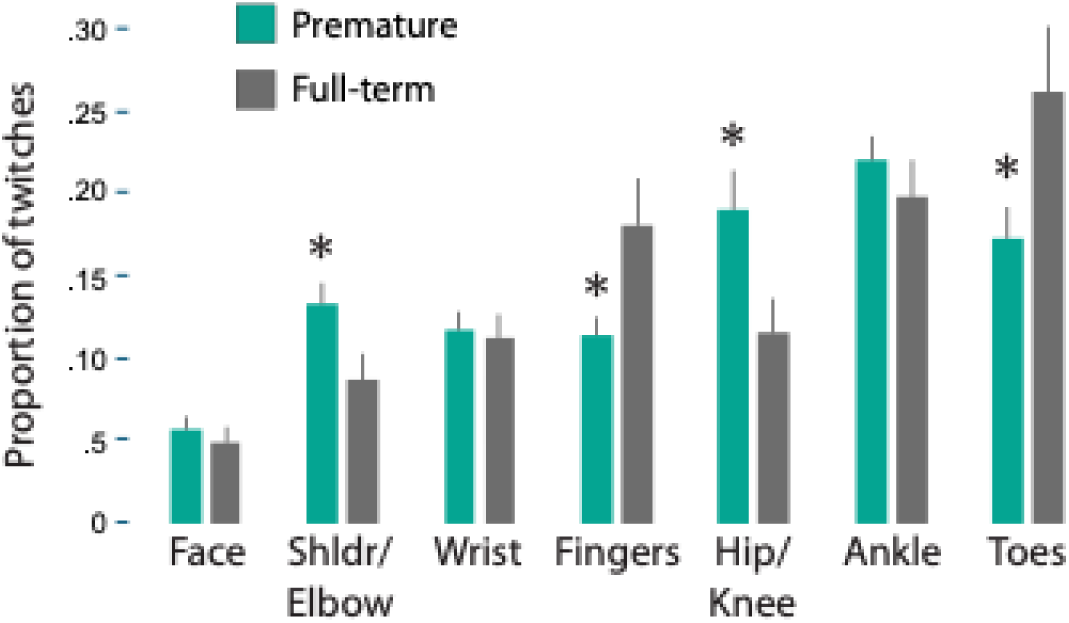
Proportion of twitches during sleep across body parts for premature and full-term infants. Data are identical to those in Figure 2c except that the data from the extremely premature and moderately premature infants are combined into a single group. * significant difference from the other group.

## Videos legends

**Video S1.** A video segment from the same moderately premature infant whose representative data are presented in Figure 1. The segment comprises a long period of AS followed by a transition to wake. Many twitches occur in the arms, legs, and face along with rapid eye movements and irregular breathing, all hallmarks of AS. The video is presented at 4x normal playback speed to make it easier to detect the movements.

**Video S2.** A video segment from the same extremely premature infant whose representative data are presented in Figure 4. The segment comprises a long period of tracé alternant. During the video, the occurrence of twitching—exclusive to the legs—is indicated. In addition to the absence of arm and face twitching, note also the absence of rapid eye movements and irregular breathing.

---

**Table S1.** Age, body weight, and basic sleep-session data for the preterm and full-term infants in this study.

| **Group** |  | **Post-menstrual age at birth** | **Age at recording session** | **Chronological age at recording session** | **Body weight at birth** | **Body weight at session** | **Duration of recording session** | **Percentage of time asleep during the recording session** | **Total duration of sleep scored** |
| --- | --- | --- | --- | --- | --- | --- | --- | --- | --- |
| **Extremely preterm**<br>(N=16; 8 female) | **Mean** | 29.1 wks | 34.7 wks | 5.7 wks | 1398.1 g | 2244.7 g | 106.9 min | 40% | 30.2 min |
|  | Min | 23.6 wks | 34.1 wks | 2.8 wks | 575 g | 1880 g | 34.0 min | 15% | 8.3 min |
|  | Max | 31.6 wks | 35.4 wks | 11.7 wks | 2300 g | 2705 g | 138.2 min | 85% | 42.9 min |
| **Moderately preterm**<br>(N=13; 6 female) | **Mean** | 33.7 wks | 34.8 wks | 1.1 wks | 2350.8 g | 2338.5 g | 104.6 min | 57% | 31.5 min |
|  | Min | 32.3 wks | 34.1 wks | 0.4 wks | 1770 g | 1920 g | 57.3 min | 11% | 5.7 min |
|  | Max | 34.4 wks | 35.7 wks | 2.7 wks | 3580 g | 3580 g | 128.6 min | 82% | 47.8 min |
| **Full-term**<br>(N=12; 6 female) | **Mean** | NA | 46.0 wks | 6.0 wks | NA | 4687.1 g | 50.6 min | 64% | 31.0 min |
|  | Min |  | 43.0 wks | 3.0 wks |  | 3550 g | 24.3 min | 28% | 10.9 min |
|  | Max |  | 48.4 wks | 8.4 wks |  | 6220 g | 90.7 min | 98% | 60.9 min |

NA: data not available.

## Acknowledgments

Research supported by a grant from the National Institute of Child Health and Human Development (R01-HD104616). The authors thank Dr. Patrick McNamara and the team of clinical research nurses in the Department of Pediatrics in the Carver College of Medicine at the University of Iowa.

## Author Contributions

G.S. and M.S.B designed the study with input from the other authors. B.J.C., T.T.C. and D.R.R. facilitated recruitment of and data collection from premature infants. G.S., T.G.C., H.C.L., L.K.K., O.K.K., D.N.K., A.S., W.L., and I.K.J. collected data from full-term infants and analyzed data from all infants. G.S., T.G.C., and M.S.B. wrote the paper. All authors provided feedback and approved the final paper for publication.

## References

1. Blumberg MS, Lesku JA, Libourel P-A, Schmidt MH, Rattenborg NC. What Is REM sleep? Curr Biol. 2020;30:R38–49.

2. Robinson SR, Blumberg MS, Lane MS, Kreber LS. Spontaneous motor activity in fetal and infant rats is organized into discrete multilimb bouts. Behavioral Neuroscience [Internet]. 2000;114:328–36. Available from: http://psycnet.apa.org/psycinfo/2000-15286-010

3. Blumberg MS, Dooley JC, Tiriac A. Sleep, plasticity, and sensory neurodevelopment. Neuron. 2022;110:3230–42.

4. Sokoloff G, Hickerson MM, Wen RY, Tobias ME, McMurray B, Blumberg MS. Spatiotemporal organization of myoclonic twitching in sleeping human infants. Dev Psychobiol [Internet]. 2020;62:697–710. Available from: https://onlinelibrary.wiley.com/doi/abs/10.1002/dev.21954

5. Sokoloff G, Dooley JC, Glanz RM, Wen RY, Hickerson MM, Evans LG, et al. Twitches emerge postnatally during quiet sleep in human infants and are synchronized with sleep spindles. Curr Biol [Internet]. 2021;1–12. Available from: 10.1016/j.cub.2021.05.038

6. Workman AD, Charvet CJ, Clancy B, Darlington RB, Finlay BL. Modeling transformations of neurodevelopmental sequences across mammalian species. J Neurosci [Internet]. 2013;33:7368–83. Available from: http://www.jneurosci.org/cgi/doi/10.1523/JNEUROSCI.5746-12.2013

7. Werth J, Atallah L, Andriessen P, Long X, Zwartkruis-Pelgrim E, Aarts RM. Unobtrusive sleep state measurements in preterm infants - A review. Sleep Med Rev [Internet]. 2017;32:109–22. Available from: 10.1016/j.smrv.2016.03.005

8. Curzi-Dascalova L, Peirano P, Morel-Kahn F. Development of sleep states in normal premature and full-term newborns. Dev Psychobiol. 1988;21:431–44.

9. Peirano P, Algarín C, Uauy R. Sleep-wake states and their regulatory mechanisms throughout early human development. J Pediatr. 2003;143:70–9.

10. Holditch-Davis D, Scher M, Schwartz T, Barr DH. Sleeping and waking state development in preterm infants. Early Hum Dev [Internet]. 2004;80:43–64. Available from: http://www.sciencedirect.com/science/article/pii/S0378378204000817

11. Reissland N, Francis B. The quality of fetal arm movements as indicators of fetal stress. Early Hum Dev. 2010;86:813–6.

12. Hof JT, Nijhuis IJM, Mulder EJH, Nijhuis JG, Narayan H, Taylor DJ, et al. Longitudinal study of fetal body movements: nomograms, intrafetal consistency, and relationship with episodes of heart rate patterns A and B. Pediatr Res [Internet]. 2002;52:568–75. Available from: http://eutils.ncbi.nlm.nih.gov/entrez/eutils/elink.fcgi?dbfrom=pubmed&id=12357052&retmode=ref&cmd=prlinks

13. Patrick J, Campbell K, Carmichael L, Probert C. Influence of maternal heart rate and gross fetal body movements on the daily pattern of fetal heart rate near term. Am J Obstet Gynecol. 1982;144:533–8.

14. Peirano P, Curzi-Dascalova L, Korn G. Influence of sleep state and age on body motility in normal premature and full-term neonates. Neuropediatrics. 1986;17:186–90.

15. Peirano P, Curzi-Dascalova L. Modulation of motor activity patterns and sleep states in low-risk prematurely born infants reaching normal term: A comparison with full-term newborns. Neuropediatrics. 1995;26:8–13.

16. Khazipov R, Sirota A, Leinekugel X, Holmes GL, Ben-Ari Y, Buzsáki G. Early motor activity drives spindle bursts in the developing somatosensory cortex. Nature. 2004;432:758–61.

17. Dooley JC, Glanz RM, Sokoloff G, Blumberg MS. Self-generated whisker movements drive state-dependent sensory Input to developing barrel cortex. Curr Biol. 2020;30:2404–2410.e4.

18. Milh M, Kaminska A, Huon C, Lapillonne A, Ben-Ari Y, Khazipov R. Rapid cortical oscillations and early motor activity in premature human neonate. Cereb Cortex [Internet]. 2007;17:1582–94. Available from: http://www.ncbi.nlm.nih.gov/entrez/query.fcgi?cmd=Retrieve&db=PubMed&dopt=Citation&list_uids=16950867

19. Whitehead K, Meek J, Fabrizi L. Developmental trajectory of movement-related cortical oscillations during active sleep in a cross-sectional cohort of pre-term and full-term human infants. Sci Rep [Internet]. 2018;8:111–8. Available from: http://www.nature.com/articles/s41598-018-35850-1

20. Colonnese MT, Phillips MA. Thalamocortical function in developing sensory circuits. Curr Opin Neurobiol [Internet]. 2018;52:72–9. Available from: https://linkinghub.elsevier.com/retrieve/pii/S0959438817302970

21. Lindemann C, Ahlbeck J, Neural SB, Hanganu-Opatz IL. Spindle activity orchestrates plasticity during development and sleep. Neural Plast [Internet]. 2016;2016:1–14. Available from: http://www.hindawi.com/journals/np/2016/5787423/

22. Molnar Z, Luhmann HJ, Kanold PO. Transient cortical circuits match spontaneous and sensory-driven activity during development. Science [Internet]. 2020;370:1–11. Available from: https://science.sciencemag.org/

23. Heck N, Golbs A, Riedemann T, Sun J-J, Lessmann V, Luhmann HJ. Activity-dependent regulation of neuronal apoptosis in neonatal mouse cerebral cortex. Cereb Cortex [Internet]. 2008;18:1335–49. Available from: http://eutils.ncbi.nlm.nih.gov/entrez/eutils/elink.fcgi?dbfrom=pubmed&id=17965127&retmode=ref&cmd=prlinks

24. Blumberg MS, Dooley JC, Sokoloff G. The developing brain revealed during sleep. Curr Opin Physiol [Internet]. 2020;15:14–22. Available from: 10.1016/j.cophys.2019.11.002

25. Rio-Bermudez CD, Blumberg MS. Active sleep promotes functional connectivity in developing sensorimotor networks. BioEssays [Internet]. 2018;304:1700234. Available from: http://doi.wiley.com/10.1002/bies.201700234

26. Bos AF, Braeckel KNJAV, Hitzert MM, Tanis JC, Roze E. Development of fine motor skills in preterm infants. Dev Medicine Child Neurology [Internet]. 2013;55:1–4. Available from: http://doi.wiley.com/10.1111/dmcn.12297

27. Ryckman J, Hilton C, Rogers C, Pineda R. Sensory processing disorder in preterm infants during early childhood and relationships to early neurobehavior. Early Hum Dev [Internet]. 2017;113:18–22. Available from: 10.1016/j.earlhumdev.2017.07.012

28. Rogers CE, Lean RE, Wheelock MD, Smyser CD. Aberrant structural and functional connectivity and neurodevelopmental impairment in preterm children. J Neurodev Disord [Internet]. 2018;10:1–13. Available from: https://jneurodevdisorders.biomedcentral.com/articles/10.1186/s11689-018-9253-x

29. Ball G, Boardman JP, Aljabar P, Pandit A, Arichi T, Merchant N, et al. The influence of preterm birth on the developing thalamocortical connectome. Cortex [Internet]. 2013;49:1711–21. Available from: https://linkinghub.elsevier.com/retrieve/pii/S0010945212002389

30. Caravale B. Cognitive development in low risk preterm infants at 3-4 years of life. Archives Dis Child - Fetal Neonatal Ed [Internet]. 2005;90:F474–9. Available from: http://fn.bmj.com/cgi/doi/10.1136/adc.2004.070284

31. Agrawal S, Rao SC, Bulsara MK, Patole SK. Prevalence of autism spectrum disorder in preterm infants: A meta-analysis. Pediatrics [Internet]. 2018;142:e20180134–16. Available from: http://pediatrics.aappublications.org/lookup/doi/10.1542/peds.2018-0134

32. Pritchard MA, Dassel T de, Beller E, Bogossian F, Johnston L, Paynter J, et al. Autism in toddlers born very preterm. Pediatrics [Internet]. 2016;137:e20151949–e20151949. Available from: http://pediatrics.aappublications.org/cgi/doi/10.1542/peds.2015-1949

33. Pharoah PO, Cooke T, Cooke RW, Rosenbloom L. Birthweight specific trends in cerebral palsy. Archives of disease in childhood [Internet]. 1990;65:602–6. Available from: http://eutils.ncbi.nlm.nih.gov/entrez/eutils/elink.fcgi?dbfrom=pubmed&id=2378516&retmode=ref&cmd=prlinks

34. Escobar GJ, Littenberg B, Petitti DB. Outcome among surviving very low birthweight infants: a meta-analysis. Arch Dis Child [Internet]. 1991;66:204–11. Available from: http://adc.bmj.com/cgi/doi/10.1136/adc.66.2.204

35. Esposito G, Paşca SP. Motor abnormalities as a putative endophenotype for Autism Spectrum Disorders. Frontiers Integr Neurosci [Internet]. 2013;7:1–5. Available from: http://journal.frontiersin.org/article/10.3389/fnint.2013.00043/abstract

36. Whyatt C, Craig C. Sensory-motor problems in Autism. Frontiers Integr Neurosci [Internet]. 2013;7:1–12. Available from: http://journal.frontiersin.org/article/10.3389/fnint.2013.00051/abstract

37. Diamond A. Close interrelation of motor development and cognitive development and of the cerebellum and prefrontal cortex. Child Dev [Internet]. 2000;71:44–56. Available from: http://pubget.com/site/paper/10836557?institution=

38. Mittal VA, Walker EF. Movement abnormalities predict conversion to Axis I psychosis among prodromal adolescents. J Abnorm Psychol [Internet]. 2007;116:796–803. Available from: http://doi.apa.org/getdoi.cfm?doi=10.1037/0021-843X.116.4.796

39. Nebel MB, Joel SE, Muschelli J, Barber AD, Caffo BS, Pekar JJ, et al. Disruption of functional organization within the primary motor cortex in children with autism. Hum Brain Mapp [Internet]. 2014;35:567–80. Available from: http://eutils.ncbi.nlm.nih.gov/entrez/eutils/elink.fcgi?dbfrom=pubmed&id=23118015&retmode=ref&cmd=prlinks

40. Arditi-Babchuk H, Feldman R, Eidelman AI. Rapid eye movement (REM) in premature neonates and developmental outcome at 6 months. Infant Behav Dev [Internet]. 2009;32:27–32. Available from: https://linkinghub.elsevier.com/retrieve/pii/S0163638308000775

41. Holditch-Davis D, Belyea M, Edwards LJ. Prediction of 3-year developmental outcomes from sleep development over the preterm period. Infant Behav Dev [Internet]. 2005;28:118–31. Available from: https://linkinghub.elsevier.com/retrieve/pii/S0163638304000888

42. Grigg-Damberger MM. The visual scoring of sleep in infants 0 to 2 months of age. J Clin Sleep Med [Internet]. 2016;12:429–45. Available from: http://jcsm.aasm.org/doi/10.5664/jcsm.5600

43. Blumberg MS, Adolph KE. Protracted development of motor cortex constrains rich interpretations of infant cognition. Trends Cogn Sci. 2023;27:233–45.

44. Blumberg MS, Adolph KE. Infant action and cognition: what’s at stake? Trends Cogn Sci. 2023;27:696–8.

45. Gassel MM, Marchiafava PL, Pompeiano O. Activity of the red nucleus during deep desynchronized sleep in unrestrained cats. Arch Ital Biol. 1965;103:369–96.

46. Gassel M, Marchiafava P, Pompeiano O. Rubrospinal influences during desynchronized sleep. Nature. 1966;209:1218–20.

47. Rio-Bermudez CD, Sokoloff G, Blumberg MS. Sensorimotor processing in the newborn rat red nucleus during active sleep. J Neurosci [Internet]. 2015;35:8322–32. Available from: http://www.jneurosci.org/cgi/doi/10.1523/JNEUROSCI.0564-15.2015

48. Hicks TP, Onodera S. The mammalian red nucleus and its role in motor systems, including the emergence of bipedalism and language. Prog Neurobiol [Internet]. 2012;96:165–75. Available from: 10.1016/j.pneurobio.2011.12.002

49. Olivares-Moreno R, Rodriguez-Moreno P, Lopez-Virgen V, Macías M, Altamira-Camacho M, Rojas-Piloni G. Corticospinal vs rubrospinal revisited: An evolutionary perspective for sensorimotor integration. Front Neurosci-switz. 2021;15:686481.

50. Basile GA, Quartu M, Bertino S, Serra MP, Boi M, Bramanti A, et al. Red nucleus structure and function: from anatomy to clinical neurosciences. Brain Struct Funct. 2021;226:69–91.

51. Blumberg MS, Seelke AMH, Lowen SB, Karlsson KÆ. Dynamics of sleep-wake cyclicity in developing rats. Proc Natl Acad Sci USA [Internet]. 2005;102:14860–4. Available from: http://www.pnas.org/content/102/41/14860.short

52. Caraux G, Pinloche S. PermutMatrix: a graphical environment to arrange gene expression profiles in optimal linear order. Bioinformatics [Internet]. 2005;21:1280–1. Available from: http://bioinformatics.oxfordjournals.org/content/21/7/1280.short

53. Dereymaeker A, Pillay K, Vervisch J, Vos MD, Huffel SV, Jansen K, et al. Review of sleep-EEG in preterm and term neonates. Early Hum Dev [Internet]. 2017;113:87–103. Available from: https://linkinghub.elsevier.com/retrieve/pii/S0378378217303250

54. O’Toole JM, Boylan GB, Lloyd RO, Goulding RM, Vanhatalo S, Stevenson NJ. Detecting bursts in the EEG of very and extremely premature infants using a multi-feature approach. Méd Eng Phys. 2017;45:42–50.

55. Christiansen TG, Sokoloff G, Long HC, Kopp OK, Karr LK, Blumberg MS. Twitches during N2 and N3 are coupled with sleep spindles but not delta oscillations in 6-month-old infants. Sleep. 2026;

56. Blumberg MS, Coleman CM, Gerth AI, McMurray B. Spatiotemporal structure of REM sleep twitching reveals developmental origins of motor synergies. Curr Biol [Internet]. 2013;23:2100–9. Available from: http://www.cell.com/current-biology/retrieve/pii/S0960982213011147

57. Blumberg MS. Homology, correspondence, and continuity across development: the case of sleep. Dev Psychobiol [Internet]. 2013;55:92–100. Available from: http://doi.wiley.com/10.1002/dev.21024

58. Petersen IT. Reexamining developmental continuity and discontinuity in the 21st century: Better aligning behaviors, functions, and mechanisms. Dev Psychol. 2024;60:1992–2007.

59. Rowe ML. A longitudinal investigation of the role of quantity and quality of child-directed speech in vocabulary development. Child Dev. 2012;83:1762–74.

60. Adolph KE, Cole WG, Komati MM, Garciaguirre JS, Badaly DD, Lingeman JM, et al. How do you learn to walk? Thousands of steps and dozens of falls per day. Psychol Sci [Internet]. 2012;23:1387–94. Available from: http://pubget.com/paper/23085640?institution=

61. Parmelee AH, Wenner W, Akiyama Y, Schultz M, Stern E. Sleep states in premature infants. Dev Medicine Child Neurology [Internet]. 1967;9:70–7. Available from: http://www.ncbi.nlm.nih.gov/entrez/query.fcgi?cmd=Retrieve&db=PubMed&dopt=Citation&list_uids=6031536

62. Fernandes M, Hanna S, Sharma A. Neurodevelopmental outcomes of extremely preterm infants: theoretical and epidemiological perspectives to guide shared-care decision-making. 2022;32:18–27. Available from: https://www.sciencedirect.com/science/article/pii/S175172222100175X

63. Kreider J, Blumberg MS. Mesopontine contribution to the expression of active “twitch” sleep in decerebrate week-old rats. Brain Res. 2000;872:149–59.

64. Pompeiano O, Satoh T. Vestibular influences on the red nucleus during sleep. Pflüger’s Arch für die gesamte Physiol des Menschen Tiere. 1967;298:159–62.

65. Krimmel SR, Laumann TO, Chauvin RJ, Hershey T, Roland JL, Shimony JS, et al. The human brainstem’s red nucleus was upgraded to support goal-directed action. Nat Commun. 2025;16:3398.

66. Gassel MM, Marchiafava PL, Pompeiano O. Phasic changes in muscular activity during desynchronized sleep in unrestrained cats. An analysis of the pattern and organization of myoclonic twitches. Archives Italiennes de Biologie [Internet]. 1964;102:449–70. Available from: http://www.ncbi.nlm.nih.gov/pubmed/14196085

67. Yamaguchi K, Goto N. Development of the human magnocellular red nucleus: A morphological study. Brain Dev. 2006;28:431–5.

68. Ulfig N, Chan WY. Differential expression of calcium-binding proteins in the red nucleus of the developing and adult human brain. Anat Embryol. 2001;203:95–108.

69. Williams PTJA, Kim S, Martin JH. Postnatal maturation of the red nucleus motor map depends on rubrospinal connections with forelimb motor pools. J Neurosci [Internet]. 2014;34:4432–41. Available from: http://www.jneurosci.org/cgi/doi/10.1523/JNEUROSCI.5332-13.2014

70. Barmack NH. Central vestibular system: vestibular nuclei and posterior cerebellum. Brain Res Bull. 2003;60:511–41.

71. Boyle R, Johanson C. Morphological properties of vestibulospinal neurons in primates. Ann N York Acad Sci. 2003;1004:183–95.

72. Houk JC, Gibson AR, Harvey CF, Kennedy PR, Kan PL van. Activity of primate magnocellular red nucleus related to hand and finger movements. Behav Brain Res. 1988;28:201–6.

73. Riddle CN, Baker SN. Manipulation of peripheral neural feedback loops alters human corticomuscular coherence. J Physiology [Internet]. 2005;566:625–39. Available from: http://doi.wiley.com/10.1113/jphysiol.2005.089607

74. Lang CE, Schieber MH. Differential impairment of individuated finger movements in humans after damage to the motor cortex or the corticospinal tract. J Neurophysiol. 2003;90:1160–70.

75. Eyre JA, Miller S, Clowry GJ, Conway EA, Watts C. Functional corticospinal projections are established prenatally in the human foetus permitting involvement in the development of spinal motor centres. Brain. 2000;123:51–64.

76. Galloway JC, Thelen E. Feet first: object exploration in young infants. Infant Behav Dev [Internet]. 2004;27:107–12. Available from: http://linkinghub.elsevier.com/retrieve/pii/S0163638303000730

77. Patel BA, Wallace IJ, Boyer DM, Granatosky MC, Larson SG, Stern JT. Distinct functional roles of primate grasping hands and feet during arboreal quadrupedal locomotion. J Hum Evol [Internet]. 2015;88:79–84. Available from: https://www.sciencedirect.com/science/article/pii/S0047248415002183

78. Almadhoob A, Ohlsson A. Sound reduction management in the neonatal intensive care unit for preterm or very low birth weight infants. Cochrane Db Syst Rev [Internet]. 2015;1:CD010333. Available from: http://doi.wiley.com/10.1002/14651858.CD010333.pub2

79. Lai TT, Bearer CF. Iatrogenic environmental hazards in the neonatal intensive care unit. Clin Perinatol [Internet]. 2008;35:163–81. Available from: http://linkinghub.elsevier.com/retrieve/pii/S009551080700084X

80. Szymczak SE, Shellhaas RA. Impact of NICU design on environmental noise. J Neonatal Nurs [Internet]. 2014;20:77–81. Available from: https://linkinghub.elsevier.com/retrieve/pii/S1355184113001002

81. Hoogen A van den, Teunis CJ, Shellhaas RA, Pillen S, Benders M, Dudink J. How to improve sleep in a neonatal intensive care unit: A systematic review. Early Hum Dev [Internet]. 2017;113:78–86. Available from: 10.1016/j.earlhumdev.2017.07.002

82. Pineda RG, Neil J, Dierker D, Smyser CD, Wallendorf M, Kidokoro H, et al. Alterations in brain structure and neurodevelopmental outcome in preterm infants hospitalized in different neonatal intensive care unit environments. J Pediatrics [Internet]. 2014;164:52–60.e2. Available from: https://linkinghub.elsevier.com/retrieve/pii/S0022347613010755

83. Mason B, Ahlers-Schmidt CR, Schunn C. Improving safe sleep environments for well newborns in the hospital setting. Clin Pediatr [Internet]. 2013;52:969–75. Available from: http://journals.sagepub.com/doi/10.1177/0009922813495954

84. Voos KC, Terreros A, Larimore P, Leick-Rude MK, Park N. Implementing safe sleep practices in a neonatal intensive care unit. J Maternal-fetal Neonatal Medicine [Internet]. 2020;00:1–4. Available from: 10.3109/14767058.2014.964679

85. Wang X, Groot ER de, Tataranno ML, Baar A van, Lammertink F, Alderliesten T, et al. Machine learning-derived active sleep as an early predictor of white matter development in preterm infants. J Neurosci. 2024;44:e1024232023.

86. Blumberg MS, Plumeau AM. A new view of “dream enactment” in REM sleep behavior disorder. Sleep Med Rev [Internet]. 2016;39:34–42. Available from: http://www.sciencedirect.com/science/article/pii/S1087079215001616

87. Grunau R, Holsti L, Whitfield M, Ling E. Are twitches, startles, and body movements pain indicators in extremely low birth weight infants? The Clinical journal of pain. 2000;16:37.

88. Datavyu: A video doding tool [Internet]. Databrary Project, New York University; 2014. Available from: http://datavyu.org

89. Dreyfus-Brisac C. Ontogenesis of sleep in human prematures after 32 weeks of conceptional age. Dev Psychobiol [Internet]. 1970;3:91–121. Available from: http://onlinelibrary.wiley.com/doi/10.1002/dev.420030203/abstract

90. Dooley JC, Sokoloff G, Blumberg MS. Movements during sleep reveal the developmental emergence of a cerebellar-dependent internal model in motor thalamus. Curr Biol [Internet]. 2021;31:5501–11. Available from: http://biorxiv.org/content/early/2021/06/25/2021.06.25.449956.abstract

91. jamovi (Version 2.6) [Computer Software]. Retrieved from https://www.jamovi.org. 2025.

92. Mordkoff JT. A simple method for removing bias from a popular measure of standardized effect size: Adjusted partial eta squared. Adv Methods Pr Psychol Sci. 2019;2:228–32.

